# Pleiotropy increases with gene age in six model multicellular eukaryotes

**DOI:** 10.1101/2024.11.19.624372

**Authors:** Reese Martin, Ann T. Tate

**Affiliations:** Department of Biological Sciences, Vanderbilt University, Nashville TN, 37235; Evolutionary Studies Initiative, Vanderbilt University, Nashville, Tennessee, USA

**Keywords:** gene duplication, genetic pleiotropy, evolutionary genomics, comparative genomics

## Abstract

Fundamental traits of genes, including function, length and GC content, all vary with gene age. Pleiotropy, where a single gene affects multiple traits, arises through selection for novel traits and is expected to be removed from the genome through subfunctionalization following duplication events. It is unclear, however, how these opposing forces shape the prevalence of pleiotropy through time. We hypothesized that the prevalence of pleiotropy would be lowest in young genes, peak in middle aged genes, and then either decrease to a middling level in ancient genes or stay near the middle-aged peak, depending on the balance between exaptation and subfunctionalization. To address this question, we have calculated gene age and pleiotropic status for several model multicellular eukaryotes, including *Homo sapiens*, *Mus musculus*, *Danio rerio*, *Drosophila melanogaster*, *Caenorhabditis elegans*, and *Arabidopsis thaliana*. Gene age was determined by finding the most distantly related species that shared an ortholog using the Open Tree of Life and the Orthologous Matrix Database (OMAdb). Pleiotropic status was determined using both protein-protein interactions (STRINGdb) and associated biological processes (Gene Ontology). We found that middle-aged and ancient genes tend to be more pleiotropic than young genes, and that this relationship holds across all species evaluated and across both modalities of measuring pleiotropy. We also found absolute differences in the degree of pleiotropy based on gene functional class, but only when looking at biological process count. From these results we propose that there is a fundamental relationship between pleiotropy and gene age and further study of this relationship may shed light on the mechanism behind the functional changes genes undergo as they age.

**Impact statement:** Pleiotropy, the phenomenon where a single gene acts on multiple traits, is fundamental to genomic organization and has profound consequences for fitness. This work identifies a previously unknown relationship between pleiotropy and gene age, highlighting the dynamism of pleiotropy across time. This relationship holds across six distantly related model organisms, suggesting that it could be a highly generalizable finding, at least among multicellular eukaryotes. Following from this work, future investigation into mechanisms dictating the prevalence of pleiotropy at the gene or cellular level could provide fundamental insight into the maintenance of pleiotropy despite the potential for constraining rapid adaptation.

## Introduction

Pleiotropy is the phenomenon in which a single gene effects multiple traits, and has wide ranging effects on development (Cheverud, 1996), genetic disease (Sivakumaran *et al*., 2011; Ittisoponpisan *et al*., 2017), signaling robustness (Guillaume & Otto, 2012; Papakostas *et al*., 2014), the evolution of new traits (Lenski *et al*., 2003; Armbruster *et al*., 2009) and organismal adaptability (Fraïsse *et al*., 2019; Hämälä *et al*., 2020; Kinsler *et al*., 2020). Pleiotropy is expected to arise in the genome as a result of exaptation, the process through which existing genes and genetic architecture are coopted for use in novel traits. The role of exaptation in promoting pleiotropy has been demonstrated theoretically using computational models of complex trait evolution (Lenski *et al*., 2003) and empirically in the flowering vine genus *Dalechampia* (Armbruster *et al*., 2009), where pollinator attractant traits are coopted for plant defense and vice versa. Pleiotropy has also been proposed to arise as a result of co-selection on traits, such as the integration between the growth of the cranial vault and facial masticatory apparatus reported in several primates (Cheverud, 1996). These mechanisms for increasing the number of pleiotropic genes and the number of traits that pleiotropic genes affect suggest that the prevalence of pleiotropy should be positively correlated with gene age. Supporting this idea in humans, older genes have elevated protein-protein interactions (PPI), a commonly used metric to determine the pleiotropic status of a gene (Yin *et al*., 2016).

The prevalence of pleiotropy does not strictly increase over time, as subfunctionalization following gene duplication events decreases the frequency of pleiotropy in the genome (Lynch & Wagner, 2008; Schmid & Sánchez-Villagra, 2010; Guillaume & Otto, 2012). Subfunctionalization in duplicated genes is predicted to arise through a process of complimentary degenerate mutations in the duplicated genes, leading to loss or degradation of multiple functions in the gene duplicates (Force *et al*., 1999) and promoting the maintenance of duplicated genes in the genome (Lynch & Force, 2000). In the case where the parent gene’s functions are completely segregated between the copies with no overlap in function, the overall prevalence of pleiotropy in the genome should be reduced as two genes are now performing the work of a single gene. Aside from subfunctionalization removing pleiotropic genes from the genome, a gene’s future adaptability is limited by the acquisition of novel traits (Fraïsse *et al*., 2019), placing a soft limit on the number of processes a gene can be involved in. These forces, acting in opposition to the constant growth in pleiotropy from exaptation and co-selection, could produce a wide range of distributions of pleiotropy across the gene age spectrum.

While we have insight into some relationships between the age of a gene and its fundamental characteristics – old genes tend to be more essential (Chen *et al*., 2012), for example, and have increased GC content and length (Yin *et al*., 2016) – we do not have a clear picture of how these different forces shake out to affect relationships between gene age and pleiotropy, in terms of both mean and variance among genes and species. For example, it is unclear if changes in the prevalence of pleiotropy over time are consistent among different metrics of estimating pleiotropy, such as PPI and biological process count. It is also unclear if the relationship between gene age and pleiotropy depends on the focal organism or if it is generalizable to large swathes of the tree of life. While previous studies have suggested that pleiotropic genes are more essential than their non-pleiotropic counterparts (Ittisoponpisan *et al*., 2017), this could be related to either the essentiality of old genes or the tendency of pleiotropic proteins to be hubs in gene signaling networks (De Bruyne *et al*., 2014; Sadhukhan *et al*., 2021). Furthermore, the exact dynamics of how the prevalence of pleiotropy changes with a gene’s age are unknown; while previous work demonstrated significant differences in PPI between old and young genes, little is known about changes on a more continuous scale (Yin *et al*., 2016).

We hypothesized that young genes would have the lowest prevalence of pleiotropy, accounting for the limited time they would have had to become involved in other traits. This expectation also aligns with previous work showing that young genes are less essential than old genes (Chen *et al*., 2012) and that pleiotropic genes tend to be highly essential. We hypothesized that middle-aged genes would be more pleiotropic than young genes, reflecting the increased time for genes to be recruited into other traits via exaptation. For the oldest genes we expected either that the prevalence of pleiotropy would continue to climb or that it would reach an equilibrium state, meaning the mean prevalence of pleiotropy in all age bins past a certain age would be equal, reflecting the net outcomes of forces adding and removing pleiotropy from the genome.

To address our hypothesis, we calculated the age and pleiotropic status of protein-coding genes from *Homo sapiens*, *Mus musculus*, *Danio rerio*, *Drosophila melanogaster*, *Caenorhabditis elegans*, and *Arabidopsis thaliana*. We selected these organisms in an attempt to cover a wide range of multicellular eukaryotic life, but were limited to model organisms with well-annotated genomes and protein resources. These organisms, while limited in scope compared to the true diversity of the tree of life, allow us to identify potentially generalizable and species-specific trends in the prevalence of pleiotropy. For a given species, a gene’s age was determined by finding the most distantly related common ancestor that shared an ortholog of that gene. Orthologs were collected from the Orthologous Matrix Database (OMAdb) (Altenhoff *et al*., 2021) and common ancestors were identified on a phylogeny generated from the Open Tree of Life (synthesis 14.8) (Redelings & Holder, 2017) that included 1900 species from the OMAdb. To evaluate the prevalence of pleiotropy in genes, we used Gene Ontology (Gene Ontology Consortium, 2021) Biological process (BP) labels for each gene, as well as protein-protein interactions (PPI) from the String DB (Szklarczyk *et al*., 2023). We elected to use both Gene Ontology Biological processes and PPI as these are complimentary measures of pleiotropy (He & Zhang, 2006). Biological processes assess the kinds of distinct actions a protein may carry out, often with different active sites, while PPI measures total interacting partners and does not capture what a protein is doing with those partners, or which active sites are involved in the interaction (He & Zhang, 2006). Our primary goal with this study is to provide a high-level but generalizable understanding of the evolution of pleiotropy, laying the groundwork for future exploration of this fundamental genomic phenomenon.

## Methods

Because the datasets for each species were retrieved from databases that used similar terms and formatting, we were able to apply the same pipelines for the analysis of each species individually. The one exception is *A. thaliana*, which lacked gene duplication data in the Ensembl database and was therefore excluded from singleton vs duplicate analysis. With that in mind, the following sections reference the *H. sapiens* dataset and its analysis but apply to all species unless otherwise noted.

For each species we retrieved the following number of genes: *Homo sapiens* (n = 19,467), *Mus musculus* (n = 21,128), *Danio rerio* (n = 27,897), *Drosophila melanogaster* (n = 13,659), *Caenorhabditis elegans* (n = 16,050), and *Arabidopsis thaliana* (n = 25,125).

### Gene age determination

The entire set of ortholog groups was downloaded from the OMA database (Altenhoff *et al*., 2021), and the full database was trimmed to only include those ortholog groups with an entry from the focal species (e.g. *H. sapiens*). From these ortholog groups, a list was generated including each unique species that were present in a *H. sapiens* ortholog group. A phylogenetic tree including only these unique species was trimmed from the Open Tree of Life. The Open Tree of Life is a purely relational tree and does not include branch length data, so all branch lengths were set to 1 during the creation of the OMA-specific phylogeny. The distance from the root of the tree to the most recent common ancestor of each species and *H. sapiens* was calculated and the age of each ortholog was set to the age of the oldest common ancestor between *H. sapiens* and the species in which the ortholog was found. For example, if an ortholog was present in several species, with the oldest one being *A. thaliana*, then the age of the ortholog would be the distance of the last common ancestor between *H. sapiens* and *A. thaliana* to the root of the tree. This means that orthologs with a lower distance from the base of the tree are older than those with a greater distance. If an ortholog could not have an age assigned to it for any reason, it was excluded from the analysis. This process removed 12 *A. thaliana* genes, 7 *D. rerio* genes, and no genes for *C. elegans, D. melanogaster,* or *M. musculus.* 455 genes were removed in *H. sapiens*; while this is noticeably more than the other species, it still left 98% of the initial human gene set intact.

### Age binning

The number of orthologs assigned to any specific age is widely variable, with older age groups, such as those genes present in the last common ancestor of all eukaryotes, comprising thousands of genes while more modern age groups, like the genes found only in primates, could have tens of genes or no genes at all. To overcome this uneven sampling across ages, orthologs were grouped into age bins with at least 1000 orthologs. This binning process started in the oldest ages and progressed by collapsing ages into a single bin until the total ortholog count exceeded 1000, then a new bin would be created and the process would repeat. Due to differences in total gene counts and the specific distributions of genes, each species has a slightly different count of total age bins.

### Determining the prevalence of pleiotropy

The prevalence of pleiotropy was determined on a per-gene basis using two methods: Gene Ontology unique biological processes (BP) and STRING database protein-protein interactions (PPI). These measures were selected because they provide complimentary insight into the pleiotropic characteristics of a gene. A gene’s BP count relates to a purely functional view of pleiotropy, where each unique function the gene carries out is counted as a BP and having a greater BP count indicates that the gene affects more traits (He & Zhang, 2006; Papakostas *et al*., 2014; Williams *et al*., 2023). A gene’s PPI count instead is agnostic to annotated functional traits and focuses instead on the number of other proteins it interacts with, which provides a different kind of proxy for the degree of pleiotropy (He & Zhang, 2006; Papakostas *et al*., 2014; Williams *et al*., 2023). BP counts were calculated by retrieving Gene Ontology data associated with that gene from the OMA database using the omadb python API (Altenhoff *et al*., 2021). All GO evidence types were used to determine total biological process involvement, but multiple entries of the same process were ignored. PPI counts were determined for each protein by counting the number of interactions in the STRING database that surpassed the .66 confidence score threshold. Confidence scores are provided for each interaction in STRING, generated using the available evidence for a protein-protein interaction, and the chosen threshold indicated high confidence that the given interaction exists (Szklarczyk *et al*., 2023). For both BP counts and PPI, larger numbers indicated more pleiotropy. At no point did we deploy a hard cutoff to determine if a gene was pleiotropic or non-pleiotropic, instead opting to derive distributions of these pleiotropic measures across ages.

### Determination of broad function

Using the Gene Ontology hierarchy of terms, we determined the broad category that each specific GO term was associated with, for example *pyrimidine nucleobase biosynthetic process* is a specific (leaf) term that falls under the *metabolic process* GO term. The broad categories into which we grouped genes were the child terms of the Biological_Process term (GO:0008150), which is the root of the biological process ontology. To each protein we then assigned association to a broad functional category if one of its functions fell in that category. Ultimately, 16 functional categories were present in each species, but for simplicity we elected to analyze only the metabolic, cellular, and developmental processes as these were among the most abundant in each species. We plotted immune system processes for the *M. musculus* and *H. sapiens* datasets as well, because the immune system is thought to be highly pleiotropic (Sivakumaran *et al*., 2011) and provided a basis of comparison to other selected processes. We plotted immune system processes only for *M. musculus* and *H. sapiens* because other species had immune system process gene counts that were too low for effective comparison.

### Duplicated gene analysis

Using the ensembl biomart (Harrison *et al*., 2024), we identified the total set of duplicated genes for *H. sapiens*, *M. musculus*, *D. rerio*, *D. melanogaster*, and *C. elegans*. *A. thaliana* was excluded from this analysis because it was not available in the biomart dataset. Each gene was then assigned a value based on the number of paralogs it had in the dataset, and any gene with at least one paralog was considered to be duplicated in further analysis.

## Results

### The prevalence of pleiotropy increases with gene age across eukaryotes

Generally speaking, genes in middle-aged and older time bins have more pleiotropy, as measured by biological process count, than genes in younger time bins (Fig. 1). This finding is also recapitulated in the protein-protein interaction (PPI) results (Fig. S1). In older age bins the distribution of biological processes tends to be unimodally distributed around the median value, but younger genes develop skewed or bimodal distribution of biological process count, with a prominent peak near the mean and a second peak around the single-process baseline. These secondary peaks are not observed in the PPI plots, where even the age bins with the lowest mean PPI are still well above the floor of 1 PPI. The statistical relationship between gene age and BP count was tested using a one-way ANOVA, and for each species there is a significant relationship between gene age and BP count (Table 1) as well as between gene age and PPI (Table S1).

**Figure 1:**
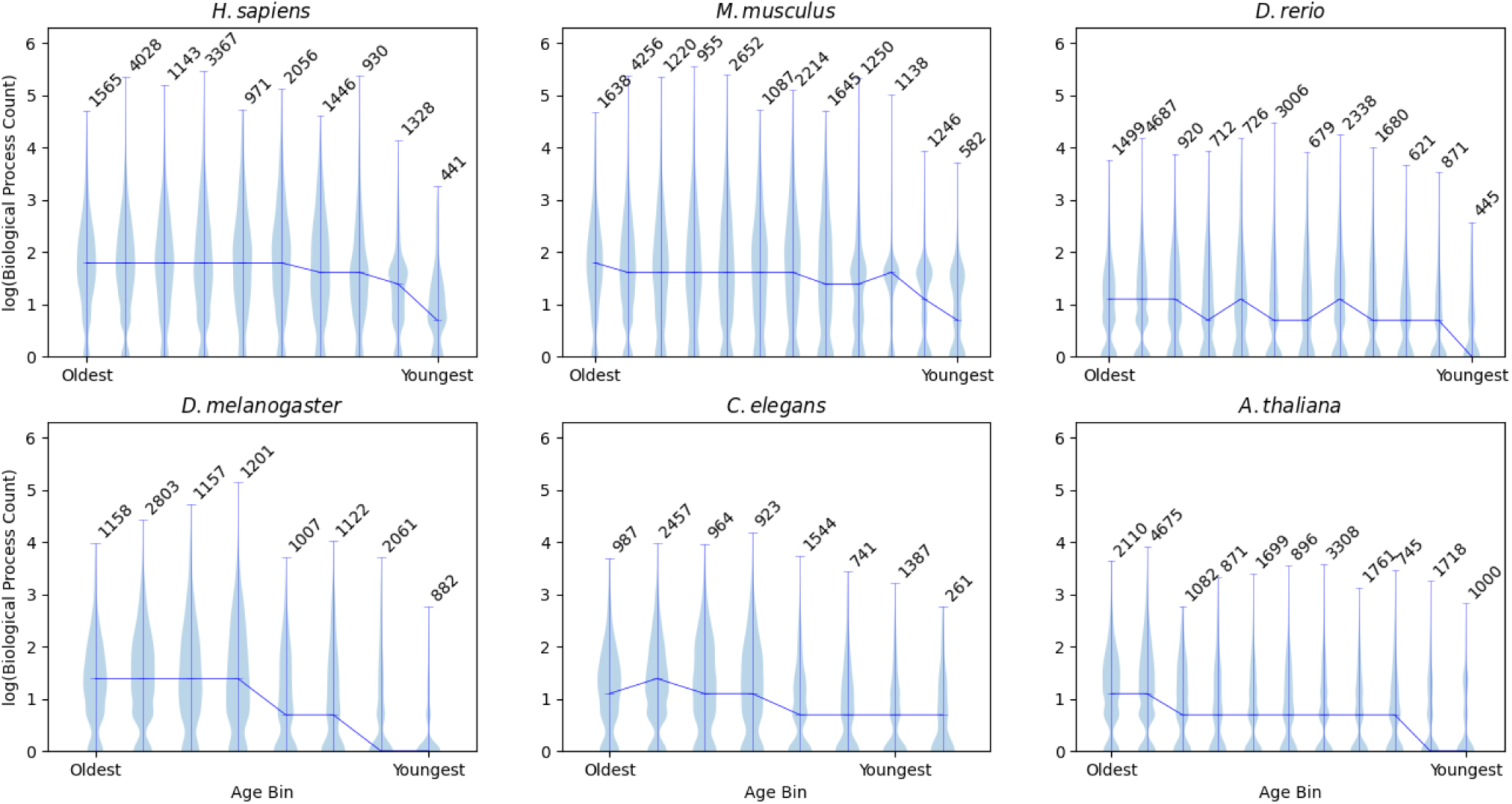
Older genes have an elevated amount of pleiotropy as measured by biological process (BP) count. The y axis shows violin plots built on the log_10_(biological process count) for all genes in each age bin. The x axis shows age bins for genes, from oldest on the left to the youngest on the right. Orthologs that did not have a BP count were excluded from these plots. The number of genes present in each binned age group is denoted above the corresponding violin.

**Table 1:** ANOVA tables for the relationship between the number of biological processes a gene is associated with and the age of that gene.

| Species | Formula | df | Sum. Sq. | Mean Sq. | F | PR(>F) |
| --- | --- | --- | --- | --- | --- | --- |
| *H. sapiens* | BP~Age | 1 | 7723.26 | 7723.26 | 49.55 | <2.01e-12 |
| *M. Musculus* | BP~Age | 1 | 2.46e4 | 2.46e4 | 163.89 | <2e-16 |
| *D. rerio* | BP~Age | 1 | 452 | 452 | 18.33 | <2e-5 |
| *D. melanogaster* | BP~Age | 1 | 2.3e4 | 2.3e4 | 521.29 | <2e-16 |
| *C. elegans* | BP~Age | 1 | 7079.67 | 7079.67 | 359.06 | <2e-16 |
| *A. thaliana* | BP~Age | 1 | 1.25e4 | 1.25e4 | 148.39 | <2e-16 |

While, broadly speaking, younger gene age corresponds to less pleiotropy than middle and older age, the qualitative dynamics of this pattern are relatively species-specific. In *H. sapiens* and *M. musculus*, the mean BP rapidly climbs as genes age and then levels out across the remaining ages. In *D. rerio* there is more variability in BP accumulation as age increases, with some age bins having lower mean BP counts than the next youngest age bin. *D. melanogaster* and *A. thaliana* exhibit a stepwise increase in BP count rather than a smooth accumulation of biological processes. *C. elegans* diverges the most from the other organisms, with its youngest age bins having approximately equal BP counts, and only the very old genes increasing in mean BP count. These patterns only manifest in the BP plots; when looking at PPI there is a universally smooth trend of constant accumulation of interactions as orthologs age.

### The prevalence of pleiotropy differs across age and functional groups

The prevalence of pleiotropy is distinct among groups of functions (Figs. 2, S2). Metabolic processes almost always have the lowest prevalence of pleiotropy, and in *H. sapiens* and *M. musculus*, the highest prevalence of pleiotropy occurs in the immune gene set in the oldest age group. For *D. rerio*, *D. melanogaster, C. elegans*, and *A. thaliana* (for which immune processes were not included) the developmental genes in the oldest age are the most pleiotropic. Interestingly these differences are not consistent between the PPI and BP plots, likely due to the kind of pleiotropy they represent (Fig. S2). Affirming our genome-wide findings, no young functional group has a higher prevalence of pleiotropy than the corresponding old functional group in the same organism. For each species, a two-way ANOVA was conducted to evaluate the relationship between age category, trait, and BP. The relationships between both age group and BP as well as trait and BP were significant for all species (Table 2). The relationship between age group and PPI was significant for all species, and the relationship between trait and PPI was significant in all species except for *D. rerio* (Table S2).

**Figure 2:**
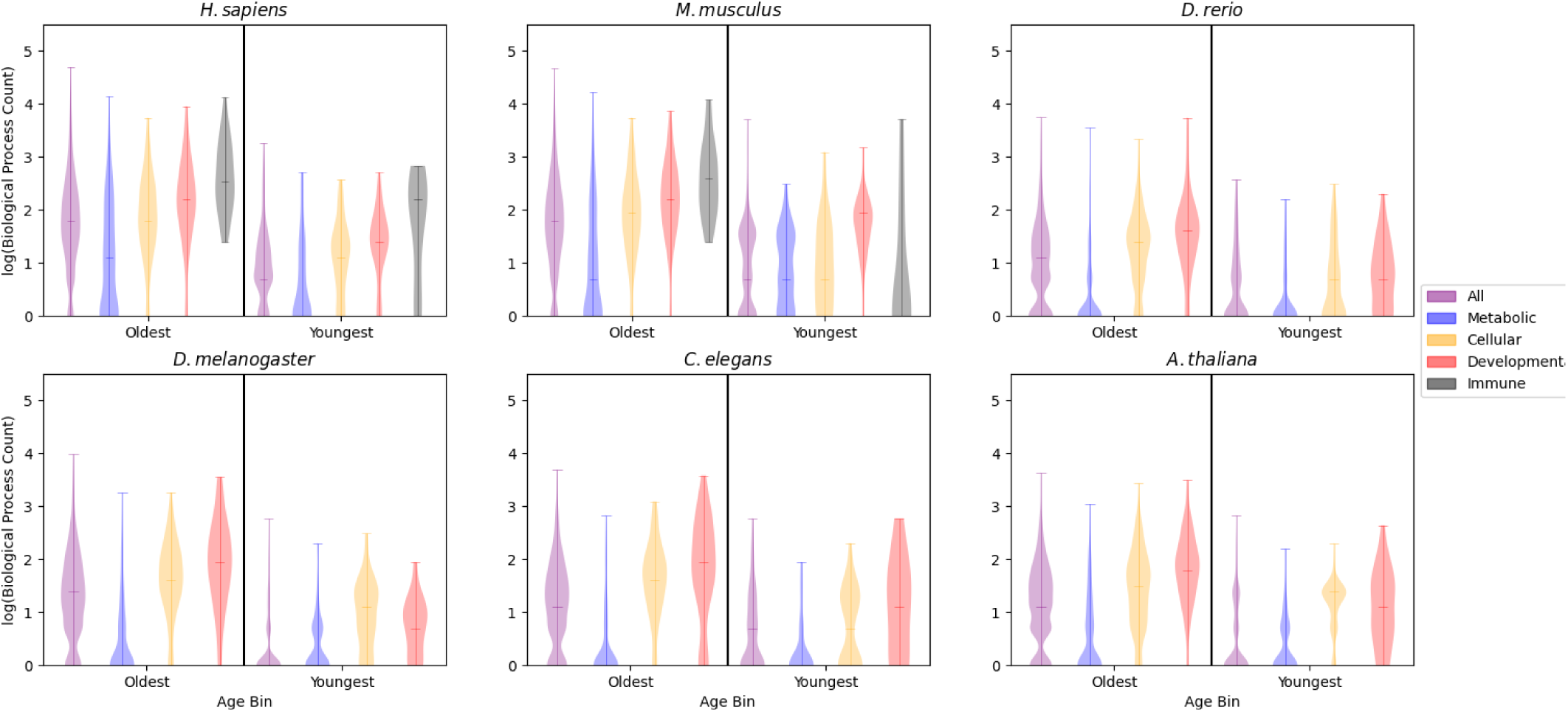
The prevalence of pleiotropy is dependent on gene function. Plots show genes in the oldest and youngest time bins (labeled Oldest and Youngest). The y axis shows violin plots built on the log_10_(biological process count) for all genes in a given age bin, separated into 4 functional groups (metabolic, cellular process, developmental, and immune processes). Orthologs that did not have a BP count were excluded from these plots.

**Table 2:** Two-way ANOVA tables for the relationship between biological process count, gene age, and the primary trait associated with a gene.

| Species | Factor | df | Sum. Sq. | Mean Sq. | F | PR(>F) |
| --- | --- | --- | --- | --- | --- | --- |
| *H. sapiens* | Age | 1 | 338.05 | 338.05 | 453.56 | <2e-16 |
|  | Traits | 4 | 166.01 | 41.5 | 55.56 | <2e-16 |
| *M. Musculus* | Age | 1 | 359.48 | 359.48 | 453.88 | <2e-16 |
|  | Traits | 4 | 174.26 | 43.56 | 55.01 | <2e-16 |
| *D. rerio* | Age | 1 | 76.46 | 76.46 | 140.26 | <2e-16 |
|  | Traits | 3 | 192.18 | 64.06 | 117.52 | <2e-16 |
| *D. melanogaster* | Age | 1 | 503.41 | 503.41 | 922.33 | <2e-16 |
|  | Traits | 3 | 150.33 | 50.11 | 91.81 | <2e-16 |
| *C. elegans* | Age | 1 | 87.75 | 87.75 | 142.13 | <2e-16 |
|  | Traits | 3 | 182.00 | 60.67 | 98.26 | <2e-16 |
| *A. thaliana* | Age | 1 | 283.68 | 283.68 | 549.60 | <2e-16 |
|  | Traits | 3 | 292.62 | 97.54 | 188.63 | <2e-16 |

The number of genes in each functional group differs significantly, so we calculated a bootstrapped mean for each functional group based on the group with the smallest number of orthologs per species. We then determined 95% confidence intervals around these bootstrapped means and found that these means were distinct based on their disjoint confidence intervals (Figs. S4, S5), suggesting that the observed differences in distributions were likely not due to sample size.

### Gene duplication events do not decrease the prevalence of pleiotropy

For each species, a two-way ANOVA was conducted to evaluate the extent to which gene age and duplication status could explain biological process count. The relationships between age and BP as well as duplication status and BP were significant for all species (Table 3). The relationship between gene age and PPI was significant for all species, and the relationship between duplication status and PPI was significant in all species except *D. melanogaster* (Table S3). Genes without paralogs generally have fewer biological processes associated with them than genes with paralogs. Several individual age bins across the species show duplicated genes with a prevalence of pleiotropy that was less than or equal to the non-duplicated genes, but the overall trend supports increased pleiotropy in genes with paralogs (Fig. 3). This trend is also seen when PPI is the measure of pleiotropy (Fig. S3). Furthermore, the exact trends between species vary, with some species showing consistent differences between duplicated and singleton genes throughout the entire age range (*H. sapiens*, *D.rerio*, *D. melanogaster*) while others have inconsistent patterns across gene age (*M. musculus*). The *C. elegans* plots are challenging to interpret due to the extremely limited duplicated gene counts.

**Table 3:** Two-way ANOVA tables for the relationship between biological process count, gene age, and gene duplication.

| Species | Factor | df | Sum. Sq. | Mean Sq. | F | PR(>F) |
| --- | --- | --- | --- | --- | --- | --- |
| *H. sapiens* | Age | 1 | 8217.78 | 8217.78 | 53.16 | <2e-16 |
|  | Duplication | 1 | 2.21e4 | 2.21e4 | 142.98 | <3.21e-13 |
| *M. Musculus* | Age | 1 | 1.64e4 | 1.64e4 | 110.97 | <2e-16 |
|  | Duplication | 1 | 4.96e4 | 4.96e4 | 334.96 | <2e-16 |
| *D. rerio* | Age | 1 | 509.25 | 509.25 | 21.00 | <4.62e-6 |
|  | Duplication | 1 | 7658.53 | 7658.53 | 315.83 | <2e-16 |
| *D. melanogaster* | Age | 1 | 2.23e4 | 2.23e4 | 517.50 | <2e-16 |
|  | Duplication | 1 | 597.49 | 597.49 | 13.85 | 1.99e-4 |
| *C. elegans* | Age | 1 | 6864.47 | 6864.47 | 348.57 | <2e-16 |
|  | Duplication | 1 | 455.62 | 455.62 | 23.14 | 1.53e-6 |

**Figure 3:**
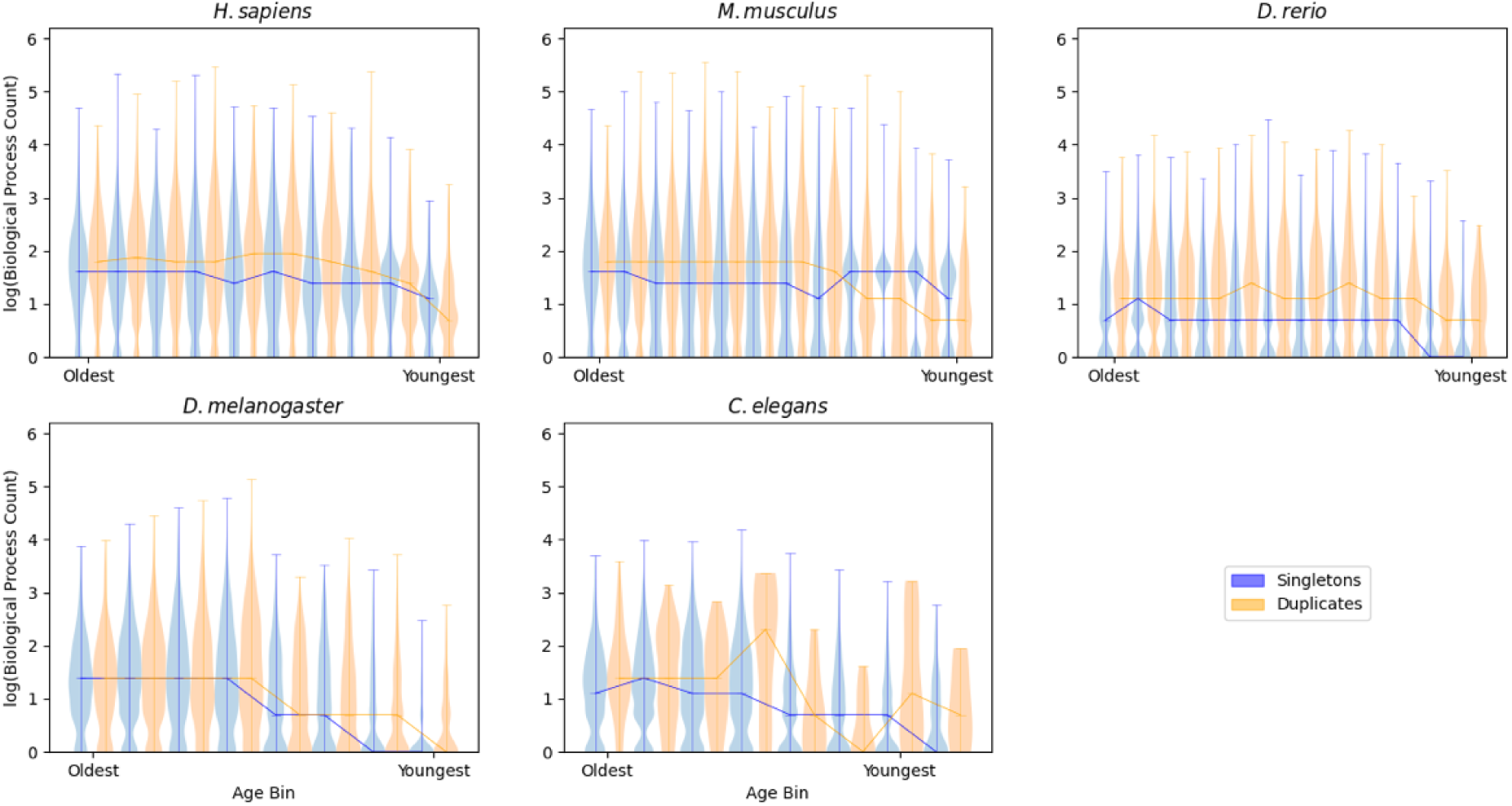
Genes with paralogs are more pleiotropic than genes without paralogs. The y axis shows violin plots built on the log_10_(biological process count) for all genes in a given age bin. The x axis shows age bins for genes, from oldest on the left to the youngest on the right. Genes with paralogs are shown in orange, genes without paralogs are shown in blue. Orthologs that did not have a BP count were excluded from these plots.

## Discussion

In this study, we have identified a potentially generalizable relationship between gene age and the prevalence of pleiotropy. Our results suggest that young genes accumulate functions as they age, eventually trending toward a plateau that could be indicative of a carrying capacity of function rather than a balance between gain and loss of function rates. We have also shown that genes belonging to metabolic, developmental, cellular, and immune functional groups differ in their prevalence of pleiotropy, and these differences are preserved within age groups. These results lay the groundwork to better understand the evolution and maintenance of pleiotropy in multicellular eukaryotes.

While previous studies suggest that both PPI and BP act as complementary proxies of pleiotropy (Williams *et al*., 2023), there are differences between the two measures that may explain the qualitative differences observed. Namely, PPI measures the number of interacting partners a protein has and is agnostic to the action of the focal protein on those partners. BP describes distinct actions a protein carries out, but any one of those processes could have many protein-protein interactions associated with it. In the most extreme cases, as with the human CAPN13 protein, a protein can be associated with a single BP (proteolysis) and have more than 750 annotated protein-protein interactions (Sorimachi *et al*., 2011). The definition of pleiotropy based on BP count is closely related to the concept of moonlighting, where a protein carries out distinct actions in different contexts (Matos *et al*., 2022). We could therefore interpret our results as indicating that gaining more interactions is easier than developing novel functionality, which matches expectations of protein evolution (i.e. micro- and macrotransitions) (Jayaraman *et al*., 2022). This explanation concurs with the observed results, where increases in PPI are largely consistent between ages (Fig. S2) while increases in BP are more abrupt and taper off significantly with age (Fig. 2).

Previous work has implicated immune systems as being highly pleiotropic (Sivakumaran *et al*., 2011; Williams *et al*., 2023), and our work has expanded on these findings by directly comparing the abundance of pleiotropy in genes across a set of traits, enabling us to determine more systemic variations in the prevalence of pleiotropy. We have shown that metabolic genes tend to have low levels of pleiotropy, as measured by PPI and BP, compared to cellular process and developmental genes. These findings are largely stable across the species we investigated (Figs. 2, S2). It is unclear if these relationships are due to differences in the ability of each trait to tolerate pleiotropy or if some traits are simply more liable to become pleiotropic through evolutionary processes. We do not expect that the differences observed are strictly due to the importance (in a fitness sense) of the traits in question as development, metabolism, and cellular processes are all fundamental to the survival and reproduction of multicellular organisms. The difference could instead be attributed to the nature of the proteins that are associated with each process. For example, signaling proteins tend to be more pleiotropic than other protein classes (Williams *et al*., 2023), potentially indicating that the abundance of signaling proteins associated with immunity and development are partially responsible for inflating the prevalence of pleiotropy in genes associated with these traits.

Despite our expectation that gene duplications would reduce the prevalence of pleiotropy, the vast majority of duplicated genes either maintain, or even increase, their prevalence of pleiotropy compared to age-matched singleton genes. A review of the functional changes associated with duplicated genes has found that complete subfunctionalization following gene duplication is likely to be rare (Janiak *et al*., 2019; Kuzmin *et al*., 2022). Several processes may act together to promote the maintenance of pleiotropy in these genes, including partial subfunctionalization where both genes maintain at least some activity for each function, neo-functionalization following subfunctionalization, dosage amplification where the excess gene products associated with multiple copies is beneficial, and backup compensation where having a second copy of a critical gene safeguards against loss of function (Kuzmin *et al*., 2022). When paralogs form complexes, selective pressure can lead to correlated mutations and provide another avenue for paralogs to actively increase pleiotropy, as one member of a complex acquiring a novel function can force others to acquire that same function (Marchant *et al*., 2019). Critically, our measures of pleiotropy cannot assess how well duplicated genes carry out their shared functions. Thus, a gene that has lost much but not all of its functional capacity following a duplication event (partial subfunctionalization) would still be pleiotropic in our analyses (Janiak *et al*., 2019). When looking for direct examples of this kind of interaction in our data we find that many duplicates, such as *myogenic differentiation 1* (*MYOD*1) and *myogenic factor 5* (*MYF*5) in mice, share a significant number of biological processes despite these genes having distinct and independent roles in specialization and differentiation (Conerly *et al*., 2016). Critically, even though *MYF5* does not induce robust transcription while *MYOD1* does, it is still associated with the biological process ‘regulation of DNA-templated transcription’. We are then left to believe that duplication events play a relatively small role in the reduction of pleiotropy in the genome, supporting the hypothesis that the accumulation of pleiotropic functions is instead limited by reduced evolvability as genes acquire novel functions (Fraïsse *et al*., 2019).

This work highlights the dynamism of pleiotropy and reveals a previously poorly understood link between a gene’s pleiotropic status and its age. This relationship holds across six distantly related model organisms, suggesting that it could be generalizable among multicellular organisms. Further work could expand these findings to the single celled eukaryotes and other domains of life to examine their broader generality. It is unlikely that the variance in the prevalence of pleiotropy observed between traits is explained by the genes in the oldest age groups being common to multiple species as we observed similar trends amongst the more species-specific young genes. However, it would be interesting to use the pseudo-replication of the same ortholog present in multiple species to study the accumulation of functions. For example, does the same ortholog have the same functions across each species? If not, are there some species where functions are similar while others have diverged? Such analysis could be expanded to conduct maximum likelihood ancestral state reconstructions to determine if the initial functions of a gene bias the other functions it may evolve over time.

Our work also suggests that the observation that immune genes are disproportionately pleiotropic in humans (Sivakumaran *et al*., 2011) may hold more broadly across species and gene age groups. This raises profound questions about the nature of genomic organization and function. For example, is the prevalence of pleiotropy dictated by the importance (in a fitness related manner) of the trait or is it intrinsic to the protein type and cellular localization that are necessary for the trait to function? Future investigation into mechanisms at the gene or cellular level could provide fundamental insight into the maintenance of pleiotropy despite the potential for constraining rapid adaptation (Guillaume & Otto, 2012; Fraïsse *et al*., 2019; Williams *et al*., 2023).

## Funding Statement

This work was supported by the National Institute of General Medical sciences at the National Institutes of Health (grant number R35GM138007 to A.T.T.).

## Author Contributions

R.A. and A.T.T. conceived the project. A.T.T. provided funding. R.A. and A.T.T. designed the analyses, and R.A. conducted them. R.A. and A.T.T. wrote the manuscript.

## Conflict of Interest Statement

The authors declare no conflicts of interest.

## Data Accessibility

The data and code used to generate these results will be made available on Dryad upon manuscript acceptance (as specified by journal policy).

## List of Supplemental Figures

**Table S1:** ANOVA tables for the relationship between the number of protein-protein interactions a gene is associated with and the age of that gene.

**Table S2:** Two-way ANOVA tables for the relationship between protein-protein interactions, gene age, and the primary trait associated with a gene.

**Table S3:** Two-way ANOVA tables for the relationship between protein-protein interactions, gene age, and gene duplication.

**Figure S1:** Old genes have an elevated prevalence of pleiotropy compared to young genes

**Figure S2:** The prevalence of pleiotropy is dependent on the functions a gene participates in

**Figure S3:** Genes with paralogs are more pleiotropic than genes without paralogs

**Figure S4:** Bootstrapped mean biological process values for functional groups of genes

**Figure S5:** Bootstrapped mean protein-protein interaction counts for functional groups of genes

## Notes

### Competing Interest Statement

The authors have declared no competing interest.

