## Supplemental Materials for "Pleiotropy increases with gene age in six model multicellular eukaryotes"

Supplemental Material for *Pleiotropy increases with gene age in six model multicellular eukaryotes*

**Table S1:** ANOVA tables for the relationship between the number of protein-protein interactions a gene is associated with and the age of that gene.

| Species | Formula | df | Sum. Sq. | Mean Sq. | F | PR(>F) |
|---|---|---|---|---|---|---|
| *H. sapiens* | PPI~Age | 1 | 2.57e6 | 2.57e6 | 240.45 | <2e-16 |
| *M. Musculus* | PPI~Age | 1 | 1.6e7 | 1.6e7 | 1218.54 | <2e-16 |
| *D. rerio* | PPI~Age | 1 | 1.31e7 | 1.31e7 | 1291.64 | <2e-16 |
| *D. melanogaster* | PPI~Age | 1 | 9.19e6 | 9.19e6 | 1422.42 | <2e-16 |
| *C. elegans* | PPI~Age | 1 | 6.53e6 | 6.53e6 | 462.06 | <2e-16 |
| *A. thaliana* | PPI~Age | 1 | 3.90e7 | 3.90e7 | 1678.95 | <2e-16 |

**Table S2:** Two-way ANOVA tables for the relationship between protein-protein interactions, gene age, and the primary trait associated with a gene.

| Species | Factor | df | Sum. Sq. | Mean Sq. | F | PR(>F) |
|---|---|---|---|---|---|---|
| *H. sapiens* | Age | 1 | 647.40 | 647.40 | 540.59 | <2e-16 |
| | Traits | 4 | 29.93 | 7.48 | 6.25 | 5.36e-5 |
| *M. Musculus* | Age | 1 | 1317.85 | 1317.85 | 1044.26 | <2e-16 |
| | Traits | 4 | 51.53 | 12.88 | 10.21 | 3.30e-8 |
| *D. rerio* | Age | 1 | 726.31 | 726.31 | 600.13 | <2e-16 |
| | Traits | 3 | 5.66 | 1.89 | 1.56 | .19 |
| *D. melanogaster* | Age | 1 | 955.98 | 955.98 | 653.70 | <2e-16 |
| | Traits | 3 | 39.71 | 13.24 | 9.05 | 5.82e-6 |
| *C. elegans* | Age | 1 | 1114.80 | 1114.80 | 591.81 | <2e-16 |
| | Traits | 3 | 25.90 | 8.63 | 4.58 | .0033 |
| *A. thaliana* | Age | 1 | 1770.34 | 1770.34 | 1303.98 | <2e-16 |
| | Traits | 3 | 16.94 | 5.65 | 4.16 | .006 |

**Table S3:** Two-way ANOVA tables for the relationship between protein-protein interactions, gene age, and gene duplication.

| Species | Factor | df | Sum. Sq. | Mean Sq. | F | PR(>F) |
|---|---|---|---|---|---|---|
| *H. sapiens* | Age | 1 | 2.83e6 | 2.83e6 | 268.22 | <2e-16 |
| | Duplication | 1 | 1.16e6 | 1.16e6 | 109.65 | <2e-16 |
| *M. Musculus* | Age | 1 | 1.58e7 | 1.58e7 | 1202.33 | <2e-16 |
| | Duplication | 1 | 2.16e5 | 2.16e5 | 16.44 | 5.04e-5 |
| *D. rerio* | Age | 1 | 1.31e7 | 1.31e7 | 1290.98 | <2e-16 |
| | Duplication | 1 | 6.33e4 | 6.33e4 | 6.23 | 0.013 |
| *D. melanogaster* | Age | 1 | 9.22e6 | 9.22e6 | 1428.41 | <2e-16 |
| | Duplication | 1 | 1.09e4 | 1.09e4 | 1.68 | 0.19 |
| *C. elegans* | Age | 1 | 6.33e6 | 6.33e6 | 448.40 | <2e-16 |
| | Duplication | 1 | 3.77e5 | 3.77e5 | 26.73 | 2.37e-7 |

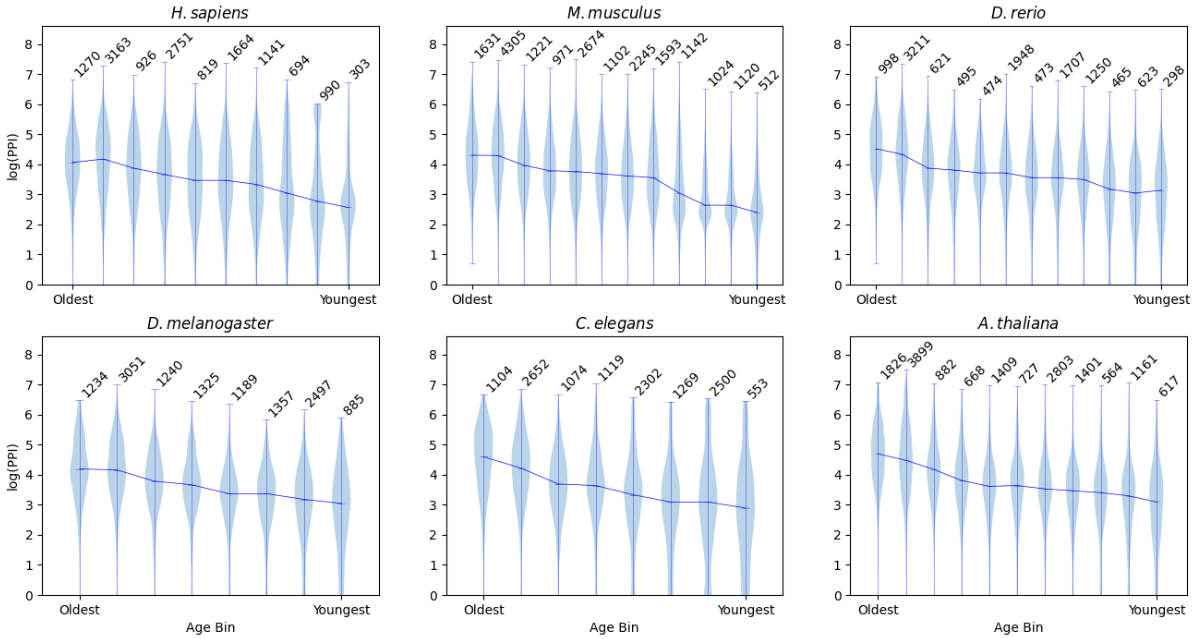


**Figure S1**: Old genes have an elevated prevalence of pleiotropy compared to young genes. The y axis shows violin plots built on the $\log_{10}$(PPI) for all genes in each age bin. The x axis shows age bins for genes, from oldest on the left to the youngest on the right. Orthologs that did not have a PPI count were excluded from these plots.

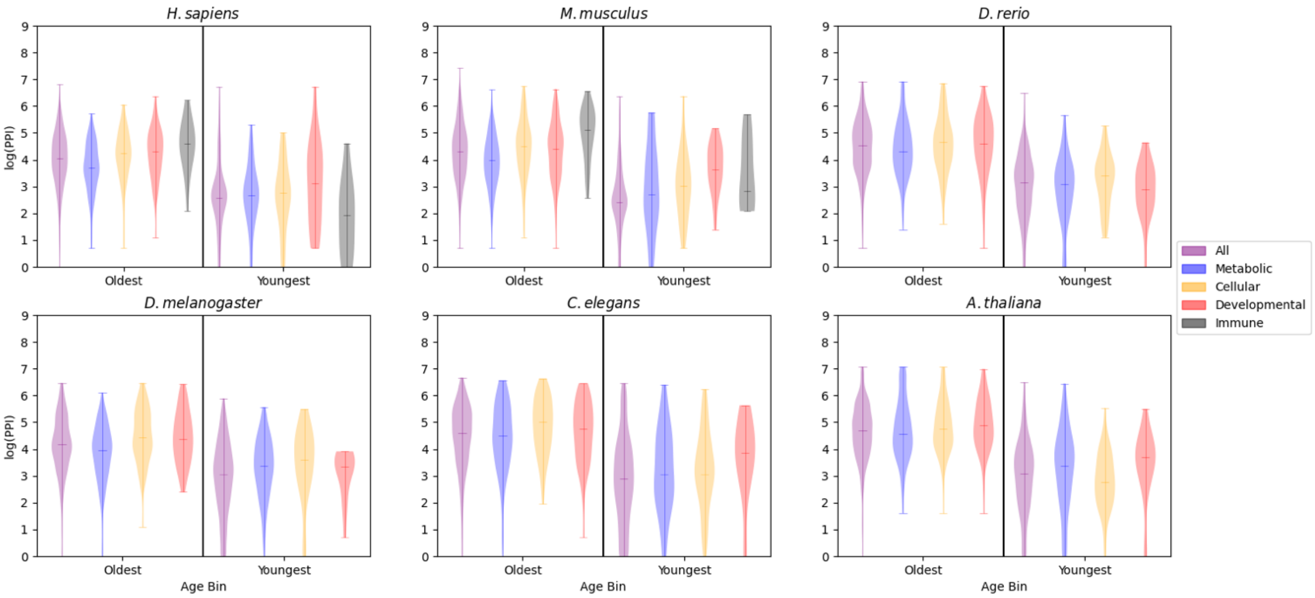


**Figure S2**: The prevalence of pleiotropy is dependent on the functions a gene participates in. Plots show genes in the oldest time bin and the youngest time bin (labeled Old and Young). The y axis shows violin plots built on the $\log_{10}$(PPI) for all genes in a given age bin, separated into 4 functional groups (metabolic, cellular process, developmental, and immune processes). Orthologs that did not have a PPI count were excluded from these plots.

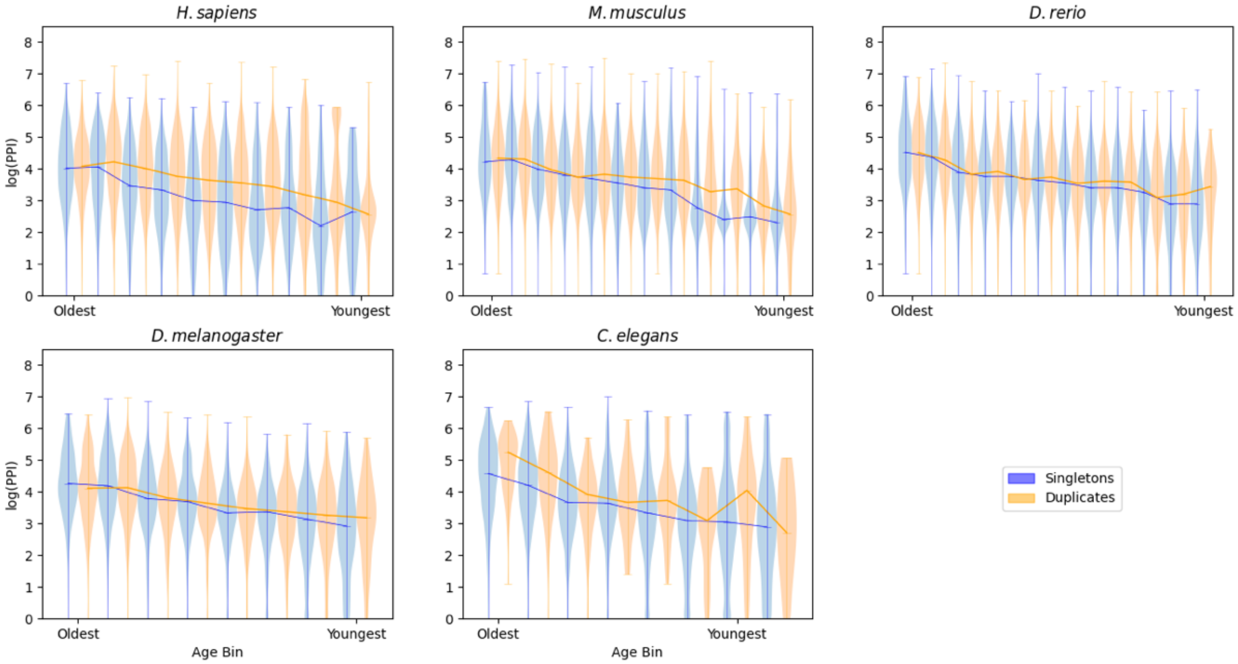


**Figure S3:** Genes with paralogs are more pleiotropic than genes without paralogs. The y axis shows violin plots built on the $\log_{10}$(PPI) for all genes in a given age bin. The x axis shows age bins for genes, from oldest on the left to the youngest on the right. Genes with paralogs are shown in orange, genes without paralogs are shown in blue. Orthologs that did not have a PPI count were excluded from these plots.

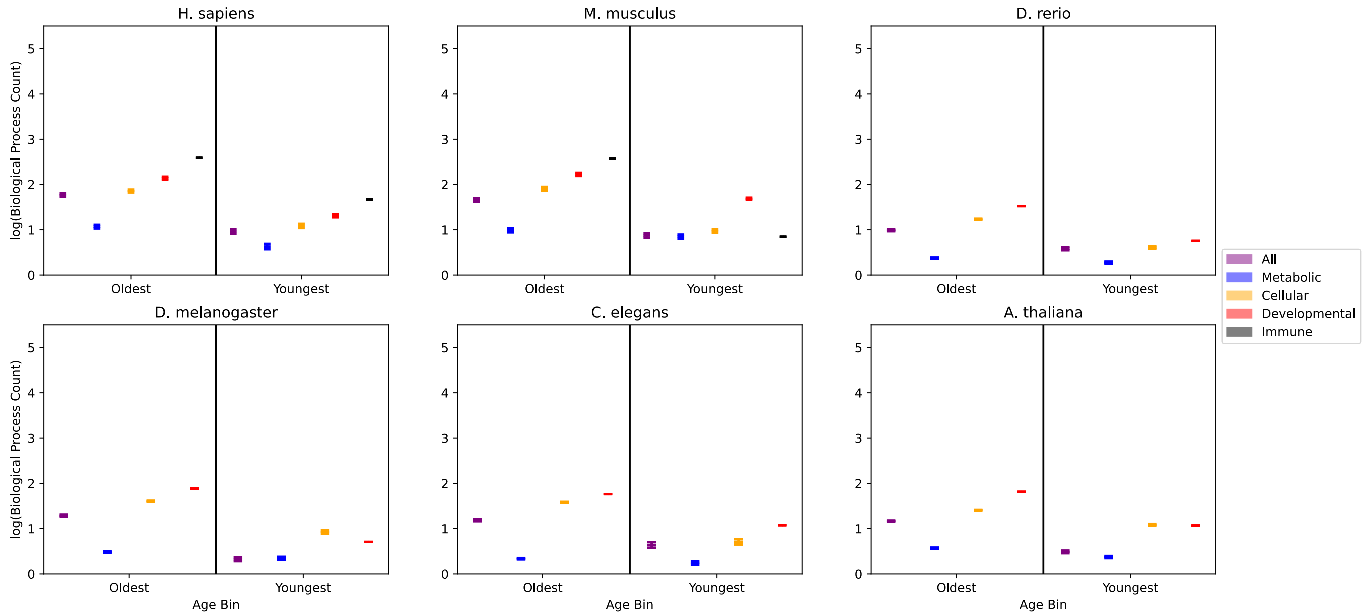


**Figure S4:** Bootstrapped mean biological process values for functional groups of genes

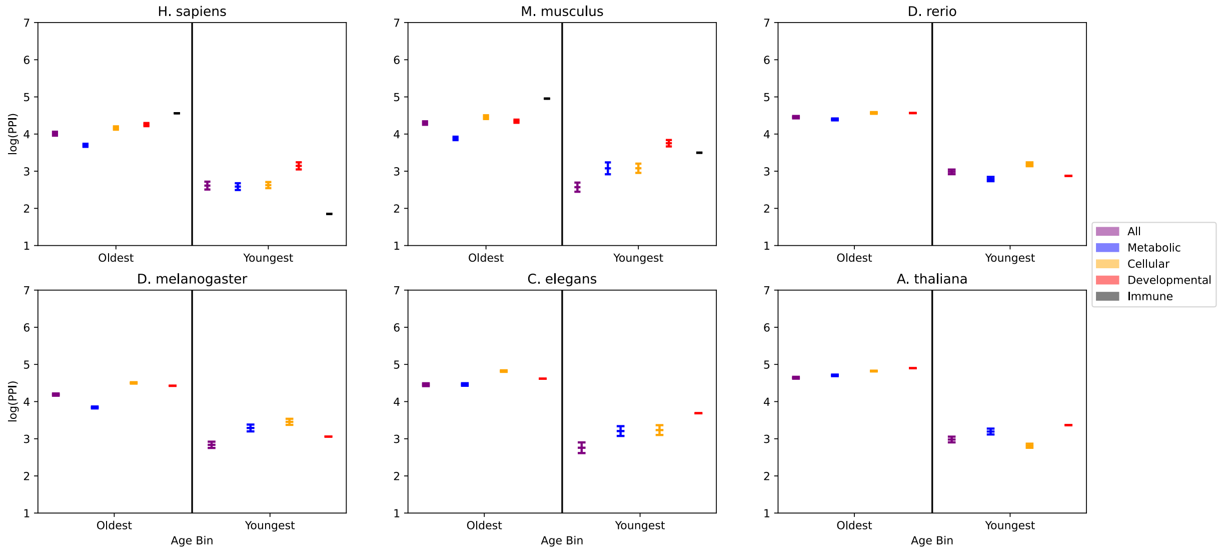


**Figure S5:** Bootstrapped mean protein-protein interaction values for functional groups of genes
